## Appendix for "Comparative Clustering (CompaCt) of eukaryote complexomes identifies novel interactions and sheds light on protein complex evolution"

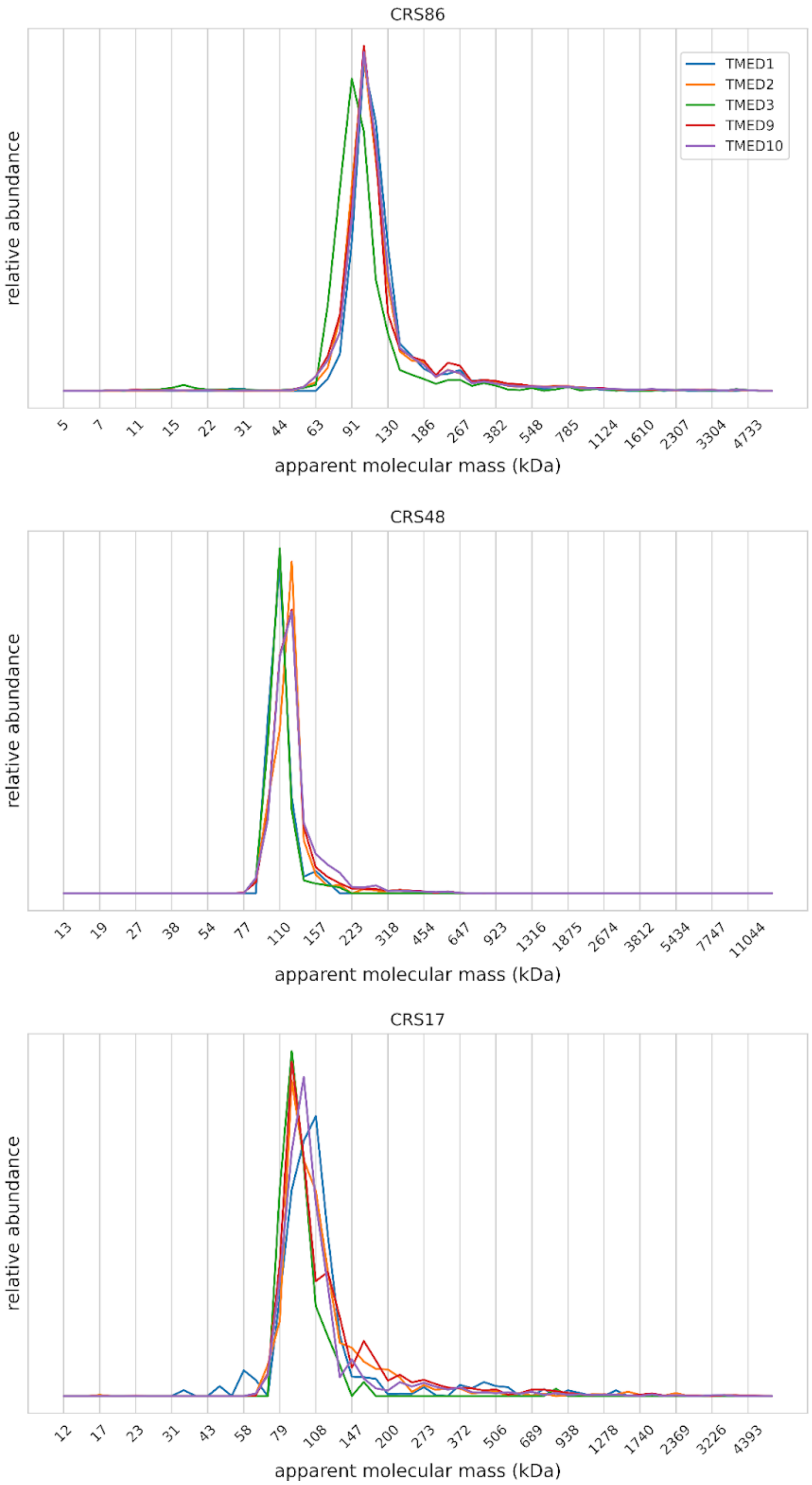


Figure S1. Migration of clustered *H.sapiens* EMP24 proteins in three human fibroblast complexome profiles. The apparent mass calibration of the complexome profiling samples was taken from the publications that first presented these data (1–3). Relative abundances were obtained by scaling the iBAQ values to a unit vector for each protein. The major peak of these TMED protein's migration patterns overlap at an apparent molecular mass of 100 to 120 kDa, corresponding approximately to the combined mass of these proteins: 125 kDa.

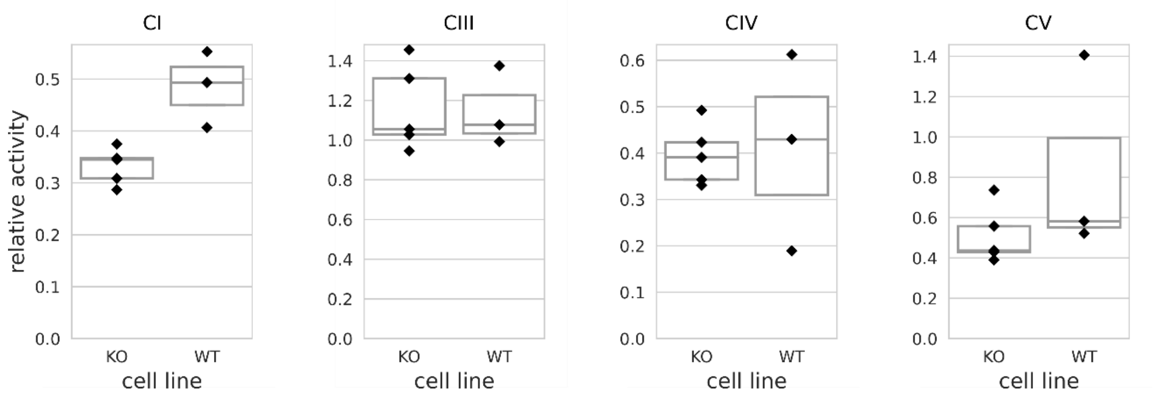


Figure S2. Enzyme activities in C15orf61-KO and WT HEK cell lines, as measured by spectrophotometric analysis. The enzyme activity values are relative to the activity of respiratory chain complex II. No significant changes were observed but complex I activities tended to be lower in the KO cell lines (Bonferroni adjusted p-values; CI: 0.092, CV: 0.583, Mann-Whitney U test). The boxes indicate quartiles and the horizontal line the median value.

```
                           cov     pid  1 [         .         .         .    ] 36
 1 Homo                 100.0% 100.0%     SFFSPGIWMGLLTSLFMLFIFTYGLHMILSLKTMDR
 2 Anopheles            100.0%  38.9%     GFTSGGILSGLFLILIFIIIGSYGIAWMMDIRTMDR
 3 Ectocarpus           100.0%  33.3%     IKITPDILAGLLTLLLFLFILVTGLGCVGDIECPKS
 4 Saccharomyces        100.0%  30.6%     SIWTEGLLMCLIVSALLLFILIVALSWISNLDITYG
 5 Naegleria            100.0%  27.8%     LMIDGEIAAGTLVGLFLLFFLVIAIVFLMNLKTSPH
 6 Arabidopsis          100.0%  25.0%     CKFKSSLLEGILVGIVFLLILISGLCCMAGIDTPTR
 7 Capsaspora           100.0%  22.2%     TWWTIPILMGLFVGAILFSILLTGVISIFTTETPTK
 8 Schizosaccharomyces  100.0%  22.2%     QFFTPGLYMGYLALAVLVPTLFISCRLLSSIQISYH
 9 Dictyostelium        100.0%  22.2%     TYVTGPVLSAYLIISILLAILFTGICCISDLQVPDR
10 Physcomitrella       100.0%  16.7%     CRSRAVILEGIFVAFTLITILVSGICCMKAVKSPAR
11 Plasmodium           100.0%  16.7%     FHTNPNILSQLMIVFLIFFFLFIGFYVLINISTPKI
12 Tetrahymena          100.0%   5.6%     QIPNQTLFGLILVFILFIPVLWIGISCLYGIESPEK
13 Toxoplasma           100.0%   2.8%     KYVTSTMLSQVVVIFLIAVTAAVGVSCLSNIDVPEI
   consensus/100%                         .h.p..hh..hhh..hhh.hh..uh..h.shp....
   consensus/90%                          .h.p.slh.thhh.hhhh.hhh.uh.hh.slp...h
   consensus/80%                          .hhs.slh.thhhhhhhh.hhhhul.hh.slpss.t
   consensus/70%                          thhssslh.slhlhhlhlhhLhhGlthl.slcsstp
```

Figure S3. Alignment of the conserved C-terminus of human ATP6AP1 and its predicted orthologs in a number of species. The *A. stephensi* (A0A182XX75) and *P. falciparum* (PF3D7_0713700) orthologs aligned here are part of the supercluster representing the $V_0$ component of the vacuolar ATPase complex resulting from our analysis. Multiple sequence alignment was performed with ClustalOmega v1.2.4 (4,5) using default settings, and the alignment was visualized with Mview v1.63 (6).

```
Reference sequence (1): sp|P24311|COX7B_HUMAN
Identities normalised by aligned length.
Colored by: identity


                                   cov    pid   1 [        .         .         .         .         :         .         .         . 80
1 sp|P24311|COX7B_HUMAN          100.0% 100.0%    -------MFPLVKSALNRLQVRSIQQTMARQSHQKR---------------------TPDF------HDKYGNAVLASGA
2 tr|Q9VVG5|Q9VVG5_DROME          80.0%  11.1%    ----------------MLVKHIVKQGLLLKNAGALSRAAYHGG-HGPHSTMNDLPVPAGDWKEQHSQKNAKYNAALITGI
3 tr|A0A182Y1F7|A0A182Y1F7_ANOST  91.2%  16.0%    LQTVQYLRFSALQETKKMQPVRSVANLAIRNAAFLSR-GYHGPSNFRVYTMNDMPVPEGDFFEEHRRKNRVYNTVLAAGI
  consensus/100%                                  .................h..hhshtthhh+put.hp.....................tsDa......+sthhNssLhsGh
  consensus/90%                                   .................h..hhshtthhh+put.hp.....................tsDa......+sthhNssLhsGh
  consensus/80%                                   .................h..hhshtthhh+put.hp.....................tsDa......+sthhNssLhsGh
  consensus/70%                                   .................h..hhshtthhh+put.hp.....................tsDa......+sthhNssLhsGh

                                   cov    pid  81          .         1         .   ] 114
1 sp|P24311|COX7B_HUMAN          100.0% 100.0%    TFCIVTWTYVATQVGIEWNLSPVGRVTPKEWRNQ
2 tr|Q9VVG5|Q9VVG5_DROME          80.0%  11.1%    LVLAGTIGFVKSSGIIHFNYYAPKSLD-------
3 tr|A0A182Y1F7|A0A182Y1F7_ANOST  91.2%  16.0%    VIFGITLTVAKESGLIYLNYNPPKSLD-------
  consensus/100%                                  hhhhhThshstppshI.hNh.sstpls.......
  consensus/90%                                   hhhhhThshstppshI.hNh.sstpls.......
  consensus/80%                                   hhhhhThshstppshI.hNh.sstpls.......
  consensus/70%                                   hhhhhThshstppshI.hNh.sstpls.......
```

Figure S4. Alignment of COX7B with recently identified *D. melanogaster* ortholog C9VVG5 and *A. stephensi* A0A182Y1F7 protein that clusters with Complex IV in our results. Multiple sequence alignment was performed with ClustalOmega v1.2.(4,5) using default settings, and the alignment was visualized with  Mview v1.63 (6).

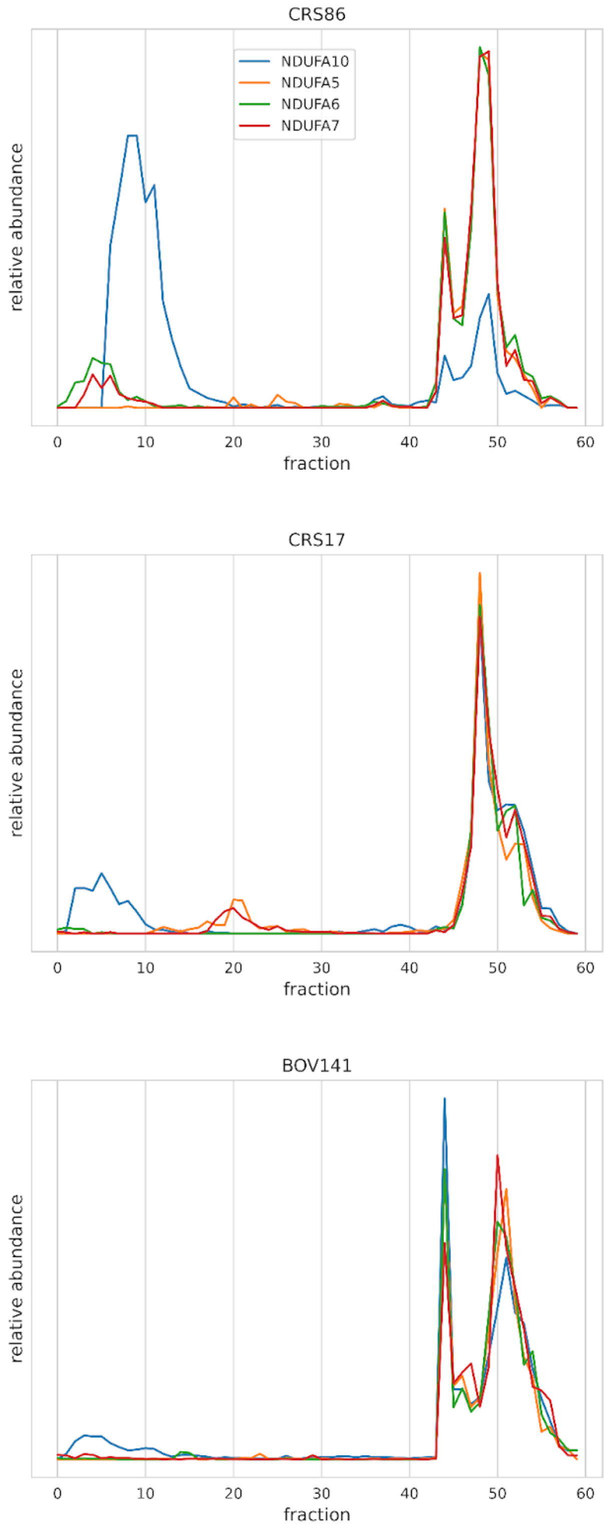


Figure S5: Migration of four complex I proteins in two *H. sapiens,* one *B. taurus* and one *A. stephensi* complexome profiles. Relative abundances were obtained by scaling the iBAQ values to a unit vector for each protein. Aside from its migration together with the other complex I subunits, it also occurs at lower abundance, suggesting that aside from being part of complex I, it is present in the samples as a monomer.

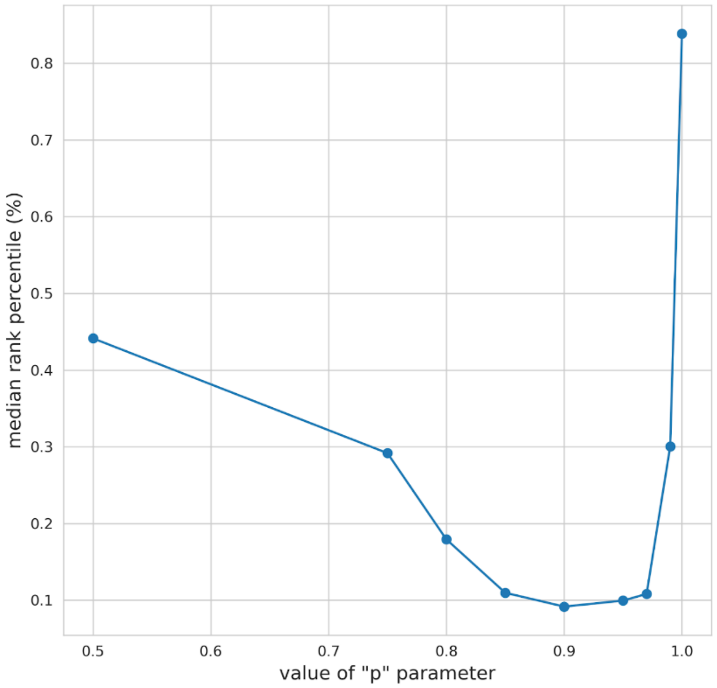


Figure S6. performance for a range of rank biased overlap (RBO) “p” parameter values for identification of proteins that are part of the same protein complex. The p parameter determines the “top heaviness” of the RBO metric, i.e., the degree to which higher ranks influence the RBO score more than lower ranks when comparing ranked lists of protein interactions. To determine the optimal RBO p parameter for the identification of pairs of proteins that are part of the same complex, RBO scores were computed for all possible protein pairs between two complexome profiles, one from *P. falciparum* and one from *T. gondii*. An evaluation set of 1285 protein pairs was created, each consisting of one *Toxoplasma* and one *Plasmodium* protein that are known to be part of the same protein complex, using a set of protein complexes conserved between these species. After ranking all protein pairs based on their RBO scores, the median rank of the evaluation set was computed, expressed in a percentile of the total list, to represent the degree to which the RBO metric scores protein pairs that are part of the same protein complex higher than other protein pairs. The median rank of the evaluation set was computed for a range of different values for “p”.

#### Table S1. Overview of 15 inspected *A. stephensi* subclusters

| Supercluster id | Complex | Predicted association | Not associated, subunit ortholog | Unknown function |
|---|---|---|---|---|
| 1 | Complex I | 30 | 2 | 1 |
| 10 | Complex II | 2 | 0 | 1 |
| 8 | Complex III | 3 | 1 | 1 |
| 9 | Complex IV | 4 | 0 | 1 |
| 3 | Complex V | 14 | 0 | 3 |
| 4 | 20S Proteasome | 6 | 0 | 0 |
| 7 | V-type ATPase V0 | 7 | 0 | 3 |
| 13 | V-type ATPase V1 | 6 | 0 | 0 |
| 15 | Prohibitin | 2 | 0 | 1 |
| 16 | Oxoisovalerate Dehydrogenase | 1 | 1 | 1 |
| 26 | Dolichyl-diphosphooligosaccharide—protein glycosyltransferase | 3 | 0 | 0 |
| 28 | Signal peptidase | 3 | 0 | 1 |
| 30 | 60S acidic ribosomal proteins | 2 | 1 | 0 |
| 31 | Electron transfer complex | 2 | 0 | 0 |
| 97 | Integrin complex | 3 | 0 | 0 |
| **Total** | 15 | 88 | 5 | 13 |

Overview of 15 inspected *A. stephensi* subclusters resulting from CompaCt analysis of eukaryote complexome profiling data, filtered using the "best guess" cluster member selection criterion. Protein members are divided in three categories based on information available in their Uniprot (31) entries: proteins that have annotations associating them with the respective protein complex, proteins whose function is unknown, and proteins that have no evidence associating them with the complex, but we predict based on sequence-based homology to be orthologous with a known subunit of the respective complex.

**Table S2. Overview of complexome profiles used in main analysis**

| **Complexome Alias** | **species** | **description** | **CEDAR experiment ID** | **samples used** | **CEDAR sample id(s)** |
|---|---|---|---|---|---|
| HUM | *H. sapiens* | fibroblast mitochondria | CRX22 | 1 | CRS86 |
| HUM | *H. sapiens* | fibroblast mitochondria | CRX17 | 1 | CRS50 |
| HUM | *H. sapiens* | fibroblast mitochondria | CRX15 | 1 | CRS48 |
| HUM | *H. sapiens* | fibroblast mitochondria | CRX9 | 4 | CRS22-25 |
| HUM | *H. sapiens* | fibroblast mitochondria | CRX8 | 1 | CRS17 |
| BOVIN | *B. taurus* | heart mitochondria | CRX33 | 6 | CRS139-144 |
| ANOST | *A. stephensi* | salivary tissue | CRX41 | 2 | CRS201-203 |
| YARLI | *Y. lipolytica* | mitochondria | CRX40 | 4 | CRS197-200 |
| PF3_GAM | *P. falciparum* | gametocyte mitochondria | CRX23 | 4 | CRS96-99 |
| PF3_AS | *P. falciparum* | asexual stage mitochondria | CRX23 | 4 | CRS92-95 |
| PF3_SCH | *P. falciparum* | blood-stage schizont whole cell | CRX20 | 6 | CRS70-75 |
| BER_SCH | *P. berghei* | blood-stage schizont whole cell | CRX20 | 6 | CRS58-63 |
| KNO_SCH | *P. knowlesi* | blood-stage schizont whole cell | CRX20 | 6 | CRS64-69 |
| TOX | *T. gondii* | tachyzoite mitochondria | CRX27 | 1 | CRS114 |
| AT_LF | *A. thaliana* | leaf mitochondria | CRX24 | 3 | CRS100-102 |
| AT_SD | *A. thaliana* | seedling mitochondria | CRX24 | 3 | CRS103-105 |

**Table S3. Overview of proteomes used for sequence-based homology analysis**

| species | proteome description | source | Reference |
|---|---|---|---|
| *A. thaliana* | Araport11_pep_20210622_representative_gene_model | https://www.arabidopsis.org | (7) |
| *H. sapiens* | UP000005640_9606 | uniprot.org | (8) |
| *B. taurus* | UP000009136_9913 | uniprot.org | (8) |
| *A. stephensi* | UP000076408_30069 | uniprot.org | (8) |
| *Y. lipolytica* | UP000001300_284591 | uniprot.org | (8) |
| *P. falciparum* | PlasmoDB-52_Pfalciparum3D7_AnnotatedProteins | https://plasmodb.org/ | (9) |
| *P. knowlesi* | PlasmoDB-55_PknowlesiH_AnnotatedProteins | https://plasmodb.org/ | (9) |
| *P. berghei* | PlasmoDB-55_PbergheiANKA_AnnotatedProteins | https://plasmodb.org/ | (9) |
| *T. gondii* | ToxoDB-52_TgondiiME49_AnnotatedProteins | https://toxodb.org/ | (9) |

### SUPPLEMENTAL METHODS

#### Generation of *P. falciparum* sporozoites / *A. stephensi* salivary gland material

*P. falciparum* strain NF54 asexual blood stage parasites were maintained in RPMI supplemented with 10% human serum and 5% haematocrit using standard culturing technique (10). To induce gametocytogenesis, asexual parasites were set at 0.5% parasitemia and left to overgrow for 14 days in a semi-automated shaker system until harvest (11). 900 µL of the gametocyte culture (approximately 2.5% haematocrit and 1-8% gametocytemia) was mixed with 540 µL packed RBCs and the cells were centrifuged for 20s at 600 x G. The supernatant was carefully removed and the pellet resuspended in 450 µL of human serum. Midi membrane feeders were used to feed female *A. stephensi* (Sind Kasur Nijmegen strain) mosquitoes (12). On day 17 after feeding, mosquitoes were dissected and the salivary glands collected and homogenized with home-made glass grinders in complete Williams B media [William's E medium with Glutamax (Thermo Fisher, 32551-087), supplemented with 1× insulin/transferrin/selenium (Thermo Fisher, 41400-045), 1 mM sodium pyruvate (Thermo Fisher, 11360-070), 1× MEM-NEAA (Thermo Fisher, 11140-035) at room temperature (13). Sporozoites were counted in a Burker-Turk chamber using phase contrast microscopy.

#### Separation of *P. falciparum* sporozoites / *A. stephensi* salivary gland material

The following steps were performed at RT with all solutions brought to RT prior to usage unless mentioned otherwise. 20% w/v Accudenz (Accurate Chemical #AN7050) was made up in demiwater and supplemented with 1× cOmplete™ EDTA-free Protease Inhibitor Cocktail (Sigma) and filter sterilized. 4.5mL were added to a conical 15mL tube. Sporozoite / salivary gland homogenate was suspended in a total of 2mL Williams B medium and carefully layered on top of the Accudenz cushion. The tube was centrifuged for 20 minutes at 2500 x G without brake. Sporozoites were extracted from top of Accudenz cushion, while salivary gland material was recovered from bottom of the tube. Both fractions were washed twice in 4°C PBS supplemented with 1× cOmplete™ EDTA-free Protease Inhibitor Cocktail and the dry pellets were flash frozen and stored at -80°C. For the non-separated sample all steps prior to washing were skipped.

#### Blue native polyacrylamide gel electrophoresis (BN-PAGE)

Protein samples were resuspended in 500 mM 6-aminohexanoic acid, 1 mM EDTA and 50 mM imidazole/HCl (pH 7.0). The samples that were separated into sporozoite and gland material were solubilized with n-dodecyl-β-D-maltoside (DDM) (Sigma) using a detergent:protein (w/w) ratio of 3:1, while the non-separated sporozoite / salivary gland homogenate was solubilized with digitonin (SERVA) at a detergent:protein (w/w) ratio of 6:1. The solubilized samples were centrifuged at 22,000 × *g* for 20 min; 4 °C. The supernatants were recovered, supplemented with Coomassie-blue loading buffer and separated on either a 4–16% or 3–16% polyacrylamide gradient blue native gels as described previously(14).

#### Cell culture conditions

HEK293T cells (ATTCC, 293T-CRL-3216) were cultured in High glucose DMEM medium supplemented with 10% fetal calf serum, 1% penicillin/streptomycin and 1% sodium pyruvate. Cells were cultured at 37 °C and 5% $CO_2$.

#### Generation of KO HEK293T cells

Two different gRNAs were designed using CRISPOR (http://crispor.tefor.net, last accessed 22 November 2022) and CHOPCHOP (https://chopchop.cbu.uib.no, last accessed 22 November 2022). Sequences of the gRNA are provided in Table S4. Two different strategies were followed: First one consisted of designing a gRNA close to the codon of the initial methionine of *C15orf61* (gRNA1). The other strategy aimed to excise a big part of the gene using two gRNAs (gRNA1 and gRNA2). gRNAs were cloned into the pSpCas9(BB)-2A-GFP (PX458) vector (Addgene, Plasmid #48138) following the protocol previously described (15). Subsequently, plasmids were transfected into HEK293T to evaluate efficacy using FuGene (Promega) reagent in a 1:3 ratio following manufacturer's instructions. After 24 h, cells fluorescent indicating that the transfection worked. Cells were harvested after 48 h and subjected to DNA isolation by incubating the pellets with MQ for 10 min at 95 °C, then adding proteinase K for 20 min at 56 °C, followed by 5 min at 95 °C to inactivate it. A standard 15 µl AmpliTaq (Life Technologies, 4398881) PCR using the primers for each of the regions (Table S5) was set as follows: 10 min at 95 °C, followed by 35 cycles of 60 sec at 95 °C, 60 sec at 54 °C and 45 sec at 75 °C, with a final elongation step of 5 min at 75°C. Subsequently, samples were loaded onto a 2%-agarose gel and were sent for Sanger sequencing to validate efficacy of gRNAs.

After validating gRNA efficacy, HEK293T at passage 20 were transfected either with gRNA1 or gRNA1+2. After 48 h, single cell sorting for GFP positive cells was performed, resulting in seeding one cell in each well of a 96 well plate. In total 2 full plates were used per condition. Clones that grew were passed to a 12 well plate, and then to a 6 well plate and a T25. Clones were sequenced for mutation identification at passage 23. Selected clones with potential promising mutations were expanded. To confirm that the editing was stable cells were sequenced at passage 25, 28 and 30. For the genotyping, for the strategy of only one gRNA, samples were amplified and sequenced using gRNA1 region primers. For the combination of two gRNAs, gRNA1 region Fw and gRNA2 region Rv were used. PCR conditions were the same as the ones described above.

*In silico* off-target analysis was performed using CRISPOR and CHOCHOP. For gRNA1 and gRNA2 no off-target sequences with an NGG PAM and no mismatches in the adjacent 12 nucleotides were found (Table S4). For gRNA1 an off-target sequence with 3 mismatches in one exon of the *UBALD1* gene was predicted by the two software's (Table S7). For gRNA2, two intergenic regions in chrosome 3 and 5 were predicted as off-target. Given the location of the possible off-targets, we validated only the *UBALD1* region as it was predicted in a gene. The region was amplified using the corresponding primers (Table S5) and by Sanger sequencing. No genome editing was observed in that region for any of the clones for which gRNA1 was used (Table S7).

For the enzyme activity data, the clones in Table S6 were used. Briefly, cells from a T75 were pelleted and frozen at passage 29.

**Table S4. gRNA sequences and off-target determined by CRISPOR and CHOCHOP**

| gRNA | Sequence 5' to 3' | PAM site | CRISPOR off-targets with 0-1-2-3 mismatches* | CHOCHOP off-targets with 0-1-2-3 mismatches* |
|---|---|---|---|---|
| gRNA1 | CAGGCGGAGCGCGACCTCGT | GGG | 0 - 0 - 0 - 1 | 0 - 0 - 0 - 1 |
| gRNA2 | TGTAGGTTAACCTAGTTCTA | GGG | 0 - 0 - 0 - 2 | 0 - 0 - 0 - 2 |

Using the most stringent filter (no mismatches in the 12 nucleotides adjacent to the PAM), both gRNA had no off-targets.

**Table S5 Primer sequences used to amplify and sequence *c15orf61* HEK293T KO**

| Primer | Sequence 5'to 3 |
|---|---|
| gRNA1 region Fw | CAAAACCAGCTCCTTGACGC |
| gRNA1 region Rv | CAGAAGGAGGTCCAGTGCG |
| gRNA2 region Fw | ACTCTGTTTAGCTGATGTGAACT |
| gRNA2 region Rv | TCCTGGCAGCTGAATGGTTT |
| Off target UBALD1 Fw | AGACCAACATCCCCTACAGC |

| | |
|---|---|
| Off target UBALD1 Rv | GGCTGTAGGGTAAGGAGCTT |

**Table S6. Characteristics of the different *c15orf61* HEK293T clones used in this work**

| Clone | Alias | Passage | Allele 1 | Allele 2 | gRNA used |
|---|---|---|---|---|---|
| HEK293T not transfected | FM-WT HEK | 29 | WT | WT | none |
| Clone 2E7 | FM-2-E7 | 29 | WT | WT | gRNA1 |
| Clone 4E7 | FM-4-E7 | 29 | WT | WT | gRNA1+gRNA2 |
| Clone 2D4 | FM-2-D4 | 29 | c.-8_132delins83 p.Met1? | c.-101_95delins45 p.Met1? | gRNA1 |
| Clone 2E11 | FM-2-E11 | 29 | c.-149_103delins149 p.Met1? | c.-149_103delins149 p.Met1? | gRNA1 |
| Clone 3C10 | FM-3-C10 | 29 | c.26_*243del p.(Ala11Cysfs*21) | c.26_*243del p.(Ala11Cysfs*21) | gRNA1+gRNA2 |
| Clone 4F5 | FM-4-F5 | 29 | c.-152_*267del p.Met1? | c.-152_*267del p.Met1? | gRNA1+gRNA2 |

**Table S7. Off-target sequences predicted by CRISPOR and CHOCHOP**

| gRNA | Off-target sequence | region | Location | Strand | MM | Result |
|---|---|---|---|---|---|---|
| #1 | CAGGCGGGGCTCGACCTCTT *TGG* | exon: *UBALD1* | chr16: 4660370-4660392 | - | 3 | No editing found |
| #2 | TGCAGGTTAAACTAGTTCGA *AGG* | intergenic: RP11-154H23.3-EIF4E3 | chr3: 71691699-71691721 | + | 3 | Not screened |
| #2 | TGTAGGTTAACCTCCTTCCA *TGG* | intergenic: C5orf66-C5orf66/H2AFY | chr5: 134642107-134642129 | - | 3 | Not screened |

Nucleotides in red indicate the mismatches with the original sequence. The PAM sequence is depicted in italics. MM: Mismatches
